# Astyanax mexicanus surface and cavefish chromosome-scale assemblies for trait variation discovery

**DOI:** 10.1101/2023.11.16.567450

**Authors:** Wesley C. Warren, Edward S. Rice, X Maggs, Emma Roback, Alex Keene, Fergal Martin, Denye Ogeh, Leanne Haggerty, Rachel A. Carroll, Suzanne McGaugh, Nicolas Rohner

**Author notes:** Authors for correspondence: Wesley C. Warren; Nicolas Rohner.

## Abstract

The ability of organisms to adapt to sudden extreme environmental changes produces some of the most drastic examples of rapid phenotypic evolution. The Mexican Tetra, *Astyanax mexicanus*, is abundant in the surface waters of northeastern Mexico, but repeated colonizations of cave environments have resulted in the independent evolution of troglomorphic phenotypes in several populations. Here, we present three chromosome-scale assemblies of this species, for one surface and two cave populations, enabling the first whole-genome comparisons between independently evolved cave populations to evaluate the genetic basis for the evolution of adaptation to the cave environment. Our assemblies represent the highest quality of sequence completeness with predicted protein-coding and non-coding gene metrics far surpassing prior resources and, to our knowledge, all long-read assembled teleost genomes, including zebrafish. Whole genome synteny alignments show highly conserved gene order among cave forms in contrast to a higher number of chromosomal rearrangements when compared to other phylogenetically close or distant teleost species. By phylogenetically assessing gene orthology across distant branches of amniotes, we discover gene orthogroups unique to *A. mexicanus*.

When compared to a representative surface fish genome, we find a rich amount of structural sequence diversity, defined here as the number and size of insertions and deletions as well as expanding and contracting repeats across cave forms. These new more complete genomic resources ensure higher trait resolution for comparative, functional, developmental, and genetic studies of drastic trait differences within a species.

## Introduction

Natural trait alterations in the Mexican tetra cavefish *Astyanax mexicanus*, some with extreme phenotypic consequences, have offered insight into how genetic adaptation works (Rohner 2018) (McGaugh *et al*. 2020). This species exists in two distinct forms: a traditional river-dweller and a blind, depigmented cave-dweller. Many studies have highlighted the use of the *A. mexicanus* surface fish to cave-life as an adaptation model to investigate the unique molecular mechanisms underlying various disease traits, such as obesity, sleep disorders, diabetes, heart regeneration, and many others, which has elevated its importance as an evolutionary model and led to its rapidly expanding use (Duboue *et al*. 2011; Aspiras *et al*. 2015) (Ojha and Watve 2018) (Stockdale *et al*. 2018) (BilandŽija 2019). Furthermore, we envision its wider use with the expansion of *A. mexicanus* genetic resources by taking advantage of a relatively short evolutionary time of transition, ∼150,000 years from surface to cave and clearly demarcated phenotypic differences (Herman *et al*. 2018).

High-quality and nearly complete chromosomes of many species are becoming more readily available, but challenges remain for genomes with difficult structural features, for example, the highly repetitive zebrafish genome (Chernyavskaya *et al*. 2022). Further research has refined best practices across broad phyla to resolve difficulties in building multiple genome assemblies with near gap-free representation (Jarvis *et al*. 2022). To date, few aquatic species have reached these higher levels of contiguity and representation, but recently developed long-read hybrid methods improve de novo assembly (Rautiainen *et al*. 2023). Many recently published aquatic genomes, although significantly more complete in sequence representation, have relied on error-prone long-read reads and their correction with short-reads (Moore *et al*. 2023) (Roberts *et al*. 2023). This new generation of genome assemblies has significantly advanced our knowledge of historically underrepresented sequences, in particular sex chromosomes (Imarazene *et al*. 2021) (Du *et al*. 2022).

In cavefish, a recent long-read based assembly improved identification of candidate genes underlying QTLs, discovered fixed or variable deletions in each cave form, and guided gene editing experiments (Warren *et al*. 2021). However, this assembly of a surface-dwelling individual suffered from a high level of gene fragmentation due to the use of high-error long reads. An individual from the Pachón cave-dwelling population was recently assembled with high levels of contiguity and facilitated the discovery of novel sex chromosome origins (Imarazene *et al*. 2021). However, given the high phenotypic divergence of the various populations, references of multiple populations are necessary to best perform comparative genomics between surface and cave morphs. To this end, we present here three highly continuous chromosome-scale assemblies of individuals from the Río Choy surface population, as well as the Molino and Tinaja cave populations to complement Pachón and for the first time present more complete references that span demographically unique and independent cave populations. The generation of these resources in *A. mexicanus* will support the community’s use of this aquatic model of human disease, particularly those wishing to compare trait diversity in the well-studied zebrafish model.

## Methods

### Sequencing and assembly

The *A. mexicanus* DNA samples were obtained from female fishes of surface, Molino, and Tinaja origins (Fig. 1A) reared at the Stowers Institute aquatic facility under IACUC approved protocol (Protocol ID: 2021-126). High molecular weight DNA was isolated using the salting out method described in the 10X genomics demonstrated protocol (10X Genomics; Pleasanton, CA) from muscle tissue to generate single molecular real time (SMRT) sequences using HiFi sequence base calling output mode from the Sequel II instrument (Pacific Biosciences) according to the manufacturer’s protocols. From all SMRT sequences, total HiFi coverage ranged from 27-46x using a genome size estimate of 1.4Gb with average read lengths across all samples of 21kb (Suppl. Table 1).

**Table 1.** Representative genome assembly metrics for sequenced *A. mexicanus* genomes.

| Common name | Assembled version | Total size Mb | Total contigs | Contig N50 length Mb | Unplaced Mb |
| --- | --- | --- | --- | --- | --- |
| Surface | Astyanax mexicanus-2.0 | 1,291 | 3,030 | 1.7 | 394 |
| Río Choy Surface | AstMex3_surface | 1,373 | 123 | 47 | 51 |
| Molino | AstMex3_Molino | 1,361 | 252 | 12 | 101 |
| Tinaja | AstMex3_Tinaja | 1,410 | 292 | 26 | 78 |
| Pachón | AMEX_1.1 | 1,378 | 529 | 14.8 | 29 |

**Figure 1.**
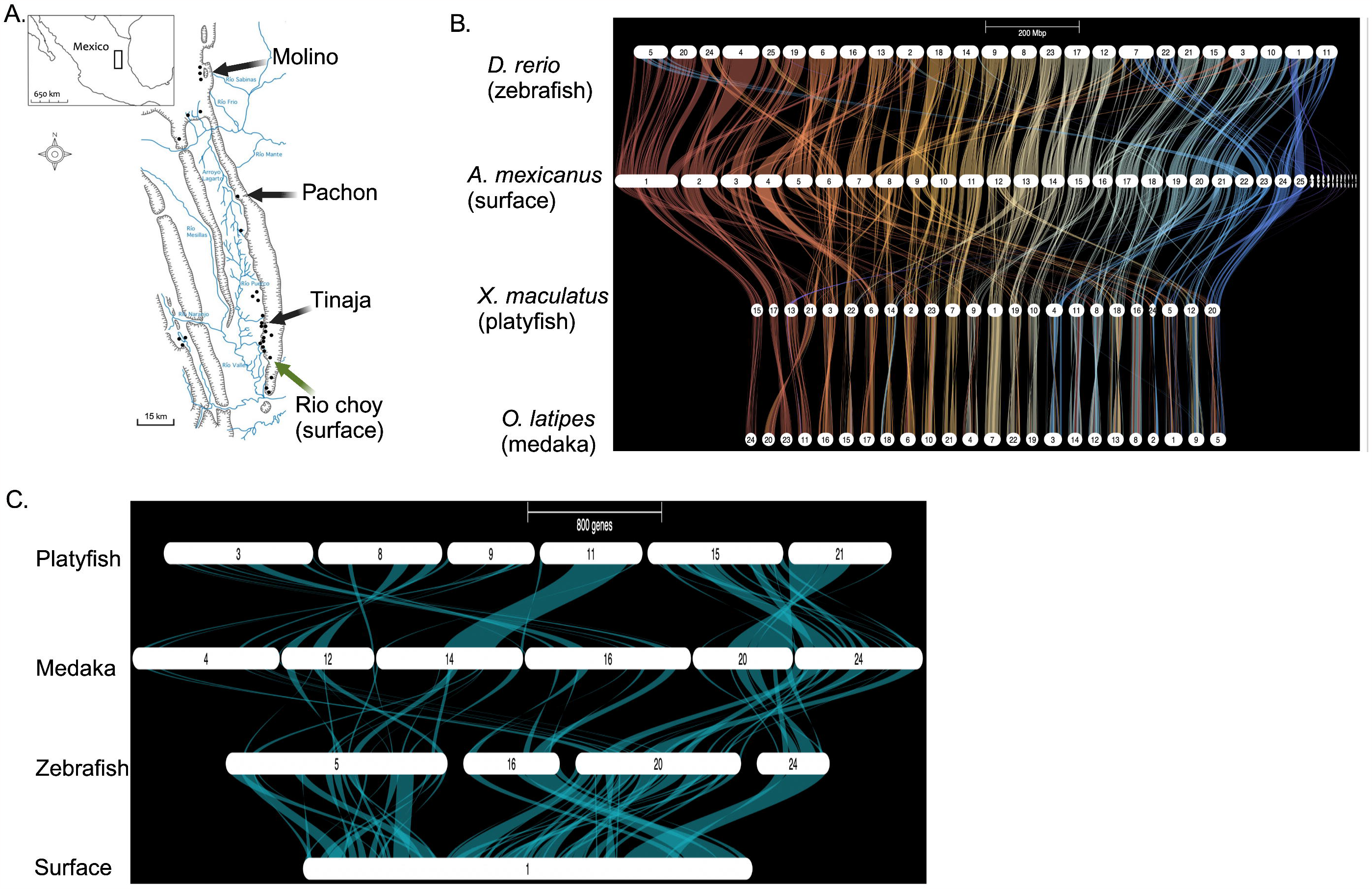
Summary of interspecies chromosomal synteny. A. Geographical representation of the independent cave morph populations and their physical appearance used in this study. Image courtesy of Alex Keene. B. Gene ortholog synteny of *A. mexicanus* against three teleost species. C. Higher resolution image of *A. mexicanus* (surface genome) aligned to the same three teleost species in B.

We assembled these reads into contigs using hifiasm v0.13-r308 with default options (Cheng *et al*. 2021). To look for haplotigs, we aligned CLR (Circular Long Reads) reads to the contigs using minimap v2.20 with arguments “-ax map-pb”. We ran purge_haplotigs v1.1.1 (Guan *et al*. 2020) on all three assemblies; the Molino and Tinaja assemblies had detectable signals of haplotigs but surface did not, so we removed haplotigs from Molino and Tinaja using coverage cutoffs (100, 197, 250) and (30, 193, 250), respectively. Haplotigs represent assembled sequences that are essentially a copy of the other haplotype and thus are removed to avoid redundancy in genome representation. To perform scaffolding a sibling of the reference fish was used to generate and sequence a HiC library developed with the Proximo Hi-C kit (Phase Genomics: Seattle, WA) according to the manufacturers protocol. We next assembled the contigs into scaffolds using a custom pipeline (see Code Availability) that aligns the Hi-C reads to the contigs with bwa mem v0.7.17 (Li and Durbin 2009), postprocesses the alignments, and finally scaffolds with SALSA v2.2 (Ghurye *et al*. 2019) and Juicebox (version 1.11.08) (Durand *et al*. 2016), as previously described (Ghurye and Pop 2019). We curated the scaffolds using a combination of synteny and manual examination of the Hi-C heatmaps (Robinson *et al*. 2018; Rhie *et al*. 2021; Warrenlab. 2022). Agptools was used to finalize chromosome assignments to be consistent with chromosome nomenclature of the surface assembled genome reported in Warren et al (Warren *et al*. 2021). Chromosomes were numbered by aligned synteny to assembled chromosomes of the previously assembled surface form of *A. mexicanus* (Warren *et al*. 2021). The completeness of gene representation was assessed using BUSCO v4.1.2 (Actinopterygii_odb10) (Simao *et al*. 2015).

### Whole genome interspecies and intraspecies analysis

To better understand shared gene organization with other teleost experimental models and across cave morphs, we first built and visualized syntenic orthology of surface fish chromosomes (AstMex3) to widely used fish experimental models including zebrafish (GRCz11), medaka (ASM223467v1), and platyfish (X_maculatus-5.0) chromosomes using the Genespace R package (Lovell *et al*. 2022). Using the parse annotation function of Genespace we generated the gene bed file from the gff files of each species. After files were parsed, formatted correctly and all headers matched in the protein and gene bed file, we initiated Genespace as described in Lovell (Lovell *et al*. 2022). We repeated this same Genespace workflow per morph compared to our surface fish assembly to evaluate possible assembly errors or natural structural variation (SV).

### Gene annotation

Gene annotation of all of our assemblies, in addition to the previously assembled Pachón (Imarazene *et al*. 2021), was carried out by the use of the standardized Ensembl workflows (Aken *et al*. 2016). However, only the surface genome described in this study was also annotated with the NCBI pipeline (Pruitt *et al*. 2014). Numerous RNAseq data sets that exist in the NCBI sequence archive for surface and cavefish from different tissue sources were used to improve the accuracy of protein-coding and non-coding gene model builds. A full accounting of all gene annotations was generated using AGAT v1.0.0 (J. 2020). We used standard parsing tools to compare lists of gene symbols in all four annotations to one another. Here, we assumed homology of matching gene symbols and ignored duplicates.

### Gene orthology

To compare gene orthology with other vertebrates, we ran OrthoFinder version 2.5.4 (Emms and Kelly 2019) on fifteen species total (Table 2). To mitigate the effect of multiple transcripts per gene, we used primary transcripts (the longest version) only (Suppl. Table 2), per OrthoFinder recommendations (Emms and Kelly 2019). MAFFT was used for multiple sequence alignment and Fasttree for gene tree inference in the OrthoFinder analysis. We ran a secondary analysis for comparison using IQtree for gene tree inference. We generated orthogroups across all species which included protein sequences from primary transcripts of *A. mexicanus* surface genome, zebrafish (*Danio rerio*), Japanese medaka (*Oryzias latipes*), platyfish (*Xiphophorus maculatus*), spotted gar (*Lepisosteus oculatus*), eastern brown snake (*Pseudonaja textilis*), common wall lizard (*Podarcis muralis*), chicken (*Gallus gallus*), human (*Homo sapiens*), chimpanzee (*Pan troglodytes*), mouse (*Mus musculus*), rat (*Rattus norvegicus*), gray short-tailed opossum (*Monodelphis domestica*), western clawed frog (*Xenopus tropicalis*), and a tunicate, *Ciona intestinalis*, as the outgroup. These taxa were chosen 1) because they all have available Ensembl annotations, thus easily compatible with Orthofinder algorithms, and 2) establish an even phylogenetic sampling across amniotes and anamniotes. In addition to the four teleosts, we include the spotted gar because this species originated prior to the teleost whole genome duplication, making it an important bridge for ortholog predictions between teleosts and other vertebrates (Braasch *et al*. 2016). We collated orthogroups that contain *A. mexicanus* sequences lacking gene symbols, so that gene symbol annotation can be inferred from that of other members in the orthogroup.

### Structural variation analysis

To initially estimate the distribution and number of SVs present in these independently derived cave morph genomes we used Assemblytics 1.2.1 (Nattestad and Schatz 2016), that estimates tandem or repeat sequence contractions and expansions, as well as deletions or insertions when compared to the surface genome (Nattestad and Schatz 2016). This approach can reflect uniqueness to a cave morph or similarity across morphs when compared to the surface genome. The nucmer algorithm of MUMmer release 4.x (Kurtz *et al*. 2004) was run to align assemblies at their contig level to minimize false positives (Nattestad and Schatz 2016). The resulting delta file was input into the Assemblytics browser and run using default parameters: a unique contig sequence anchoring size of 10,000 bp, and the variants classified by type and size ranging from 50-500 bp and 500-10,000 bp for plot visualization.

## Results and Discussion

### Chromosome-scale assembly of the surface and cavefish forms

We generated reference genomes from single lab-reared *A. mexicanus* surface or cave morphs that are descended from varying Mexico localities, representing independent origins of the cave form from Tinaja and Molino populations as well as a surface Río Choy population. We sequenced and assembled each genome using SMRT CCS (circular consensus sequencing) with a genome coverage depth of 27.4 to 46.2 (Ruan and Li 2020) (Suppl. Table 1). The final *de novo* assemblies resulted in ungapped assembly length ranges of 1.36-1.41 Gb, total contigs of 123-292, and N50 contig lengths of 12-47Mb (Table 1). Using on average 200 million 150bp Hi-C reads for chromosome-scale scaffolding resulted in the generation of 25 total chromosomes for the surface and each cave morph. Due to the exceptional contiguity of the surface assembly (47 Mb N50 contig length), we established the chromosome structure of this assembly first, then investigated discrepancies of the other cavefish assemblies using multiple pairwise alignments, including the recently published long-read assembly of the Pachón morph (Imarazene *et al*. 2021). Few order or orientation errors were found and corrected; however, three chromosomes with complete orientation differences in all cave forms must be further investigated for possible curation fixes (Suppl. Fig. 1). The surface genome contiguity surpassed all our cave morphs and, to our knowledge, all long-read assembled teleost genomes to date found in the NCBI assembly archive, including the zebrafish (fDanRer4.1). Across all assemblies, only 3.7 to 7.4% of sequences could not be properly assigned to chromosomes. These new assemblies contribute 90 Mb in new sequence on average and show a substantial reduction (25-fold) in assembly gaps, when evaluated against the prior version of the surface fish genome (A. mexicanus-2.0; Table 1). Our surface assembly exhibits a high level of contiguity and completeness, despite that the total interspersed repeats for the surface genome was 45.8%, which is only slightly lower than zebrafish’s 48.4% with a similar estimated genome size of 1.4 Gb.

### Structural differences

One question we wished to address was: is gene order highly conserved among surface and cave morph chromosomes despite their demographic origin differences (Fig. 1A) and up to 190,000 generations of cave morph divergence from their surface ancestor (Moran R.L. 2022). We find no major chromosomal discrepancies in chromosome order when performing pairwise comparisons of the Pachon, Tinaja, or Molino cave morphs to the representative surface fish ancestor (Suppl. Fig. 1). Also, of interest was: are there examples of unexpected teleost interspecies conservation in chromosome gene order that can aid future studies aimed at understanding cave morph standing genetic variation. The alignments of all 25 *A. mexicanus* chromosomes to three distantly related teleost genomes (zebrafish, platyfish, and medaka) using aligned genes are expected to display patterns of organizational divergence dependent on their phylogenetic relationships. Overall, 56% of *A. mexicanus* chromosomes display complex genomic rearrangements relative to all three distantly related teleost species (Fig. 1B). In one example, the largest *A. mexicanus* chr1 (134 Mb), aligns with four zebrafish chromosomes: 5, 16, 20, and 24 (Fig. 1C), suggesting multiple fissions or fusions occurred throughout teleost genome evolution. In contrast, there were some chromosomes that are nearly syntenic for gene order, such as *A. mexicanus* chr14 versus zebrafish chr17 (Fig. 1C). A separate pairwise alignment of the surface fish and zebrafish genomes (Danio rerio GRCz11) support these findings of variable synteny (Suppl. Fig. 2). These teleost interspecies alignments confirm earlier studies, and highlight the complex trajectory in the evolution of the teleost genome (Volff 2005) (Braasch *et al*. 2016).

**Figure 2.**
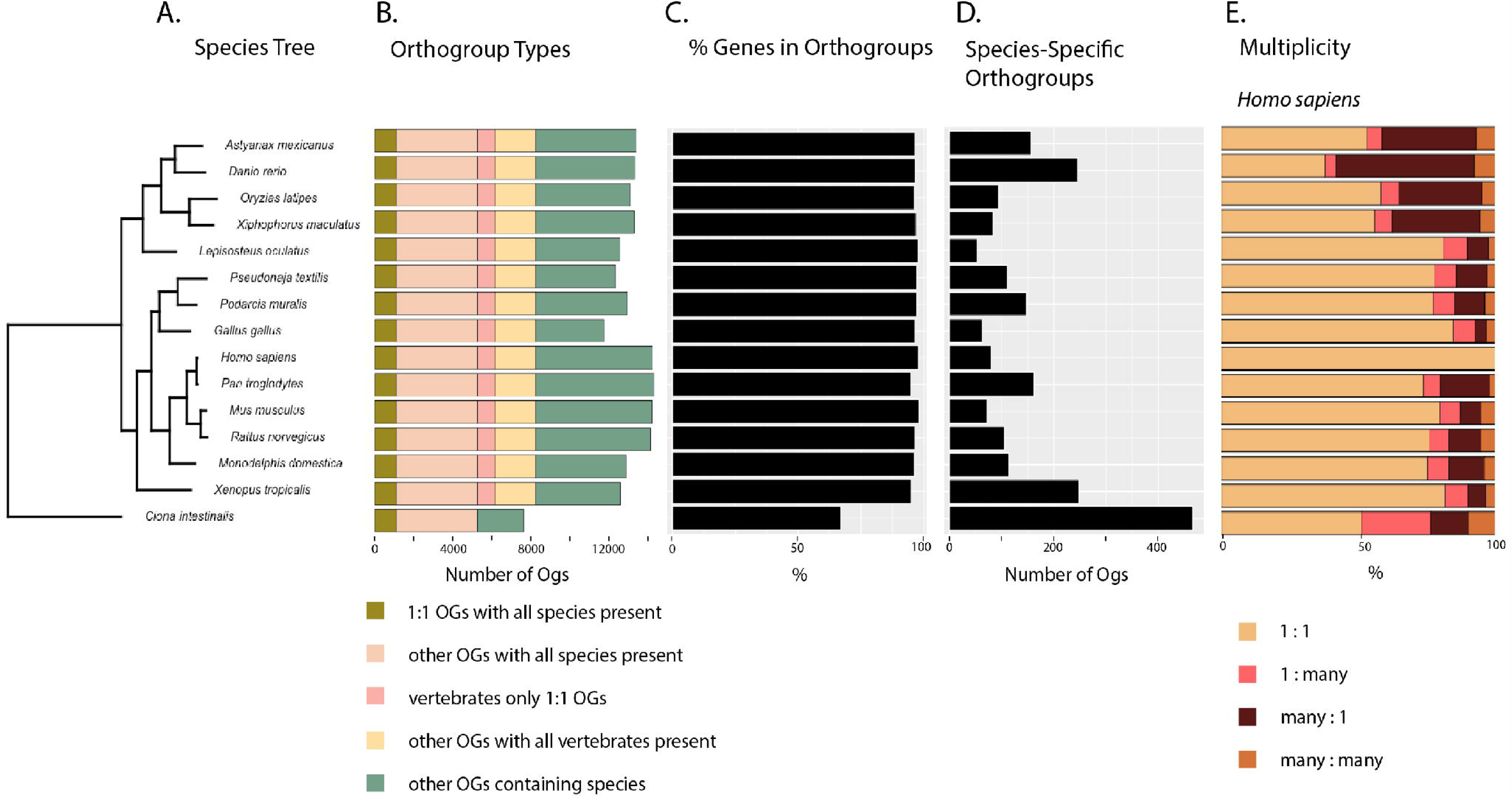
Summary of OrthoFinder analysis of vertebrate species *Astyanax mexicanus, Danio rerio, Oryzias latipes, Xiphophorus maculatus, Lepisosteus oculatus, Pseudonaja textilis, Podarcis muralis, Gallus gallus, Homo sapiens, Pan troglodytes, Mus musculus, Rattus norvegicus, Monodelphis domestica, Xenopus tropicalis*, and *Ciona intestinalis*. Bar charts describe data for each species, aligned to the matching species in the tree. A. Phylogenetic tree built with all species using shared gene orthologs. B. Number of orthogroups each classified by type per species. C. Percentage of genes by orthogroups by species. D. Number of species-specific orthogroups per species. E. Ortholog multiplicities for all species.

### Gene annotation

We first assessed the sequence completeness of the assemblies using benchmarking universal single-copy ortholog (BUSCO) (Simao *et al*. 2015) scores and found on average 96.8% of BUSCOs present in their complete and unfragmented form, 2.4% missing, and 1.3% duplicated (Suppl. Table 3). Protein-coding genes from all four genomes were predicted using Ensembl (Aken *et al*. 2016) with the average being 26,974. The surface genome was also annotated using the NCBI workflow, with small differences in the total protein-coding genes count (Suppl. Table 4). In contrast, large differences were seen in the surface genome pseudogene annotation, 1,376 versus 183, when comparing the NCBI and Ensembl output (Suppl. Table 4). Interestingly, a small increase in Ensembl predicted pseudogenes in cave morphs compared to the surface is observed (Suppl. Table 4) apart from Pachon. The correct annotation of pseudogenes “non-functional” genes across species is a persistent problem, and especially deserves further attention in cave species genomes given its importance in understanding troglomorphic genetic adaptation (Harrison 2021). While all these newly annotated *A. mexicanus* genomes are improved over prior assemblies (Warren *et al*. 2021), estimates of non-coding genes were substantially improved (a 36-fold increase; Suppl. Table 4), owing mostly to improved assembly contiguity and accuracy. In total, all these surface and cave morph gene sets show Molino, Tinaja, and Surface have similar numbers of mRNAs, exons, CDSs, and total CDS lengths, but in most comparisons, the increased surface genome contiguity resulted in improved gene annotation, supporting the use of AstMex3_surface as the standard reference for future *A. mexicanus* computational experiments (Suppl. Table 5).

In pairwise comparisons, the surface genome had 1,041, 978, and 1,054 gene symbols not found in Pachon, Molino, and Tinaja annotations, respectively (Suppl. Table 6). Similarly, each cave population had over 800 gene symbols not found in the surface genome.

### Gene Orthology

We targeted a specific span of model species across the vertebrate phylogeny to classify potential orthologs (Fig. 2A). By analyzing all detected gene ortholog relationships, we were able to better understand the utility of *A. mexicanus* for comparative inference across aquatic models and other vertebrates (Fig. 2B). In the OrthoFinder analysis, 95% of all genes were assigned to 20,612 orthogroups (Suppl. Table 7; Fig. 2C). Fifty percent of genes assigned to orthogroups are in orthogroups of 19 genes or more, and over 25% of orthogroups contained all species (n = 5,285) (Fig. 2D), of which 20.8% are single copy orthogroups (n = 1,097) (Suppl. Table 7; Fig. 2E). An additional 14.4% of orthogroups (n = 2,988) contained at least one copy from every vertebrate species and zero copies from the outgroup (Fig. 2B). All vertebrates had >96% of their genes assigned to orthogroups (Fig. 2C; Suppl. Table 8). The outgroup, *C. intestinalis*, had 66.8% of genes assigned to orthogroups (Suppl. Table 8). For *A. mexicanus*, 96.3% of genes were assigned to orthogroups (Suppl. Table 8). We found more one to one orthologs of *A. mexicanus* with humans than for zebrafish with humans (n = 6,341 and n = 5,118, respectively; Fig. 2E). We identified 378 orthogroups between *A. mexicanus* and humans that lack zebrafish orthologs (Fig. 2E). Overall, more orthogroups contain genes from *A. mexicanus* and humans (n = 11,444) than zebrafish and humans (n = 11,272). Addressing specifically orthologous relationships, 16,582 human genes (70.5%) have an ortholog with *A. mexicanus*, whereas 16,221 human genes (68.9%) have an ortholog with *Danio rerio*. These findings increase the breadth of orthologs available in an alternative fish model for comparative trait dissection. Zebrafish have more many-to-one and many-to-many orthologs with humans and more species-specific orthogroups than *A. mexicanus* (n = 245 and n = 157, respectively; Fig. 2E). In general, the availability of higher-quality genome assemblies is revealing teleost whole genome duplication to be a pervasive source of genetic variation across these taxa (Albalat and Canestro 2016), and understanding the main evolutionary forces impacting *A. mexicanus* gene losses or gains is beyond the scope of this study (Adrian-Kalchhauser *et al*. 2020). In identifying orthologs, we also estimated the number of duplications across the species tree, both at the tips and internal nodes which could represent differential genome fractionation across lineages stemming from the teleost whole genome duplication or lineage-specific duplications. At the base of teleosts, our analysis identified 2,451 duplications. In the tip branches of teleosts, we identified 2,647, 2,825, 4,812, and 8,509 duplicates for *X. maculatus, O. latipes, A. mexicanus*, and *D. rerio*, respectively. We suspect that teleost whole genome duplication, annotation bias, phylogenetic sampling, and multiple evolutionary events or forces underpin these results. Five orthogroups (OG0000022, OG0000023, OG0000045, OG0000050, and OG0000086) each have over 100 genes from *A. mexicanus*. These uniquely large copy numbers suggest that these genes experienced unique evolutionary pressures in *A. mexicanus*. The orthogroup with the largest number of *A. mexicanus* gene copies is OG0000022 (n = 215). This group and the group with the second largest number of *A. mexicanus* genes, OG0000023 (n = 205), are dominated by *A. mexicanus* zinc finger proteins. Additional studies using independent tests of gene family expansions and contractions, such as CAFE (Mendes et al. 2021), will be needed to truly resolve duplication events prior to explorations of their functional roles in the *A. mexicanus* cave adaptation. We provide gene symbols for human and *A. mexicanus* orthologs as a resource for comparative studies (Suppl. Table 9). These *A. mexicanus* protein-coding gene resources further efforts to improve genome annotation for *A. mexicanus*, offering another aquatic model beyond zebrafish for human comparative studies. Future research to identify gene duplicates, pseudogenization events, and nonsynonymous protein coding changes unique to *A. mexicanus* within a framework of surface and independent cave morphs will address hypotheses about how this species was adaptable to extreme environmental change.

### Structural variation

Cave morphs display many phenotypic differences from their surface ancestor that motivated us to evaluate genome evolution on a finer scale beyond chromosomal gene order evaluated herein and prior studies that have focused on protein-coding gene changes (Warren *et al*. 2021) (Moran R.L. 2022). Discovery of SVs among cave morphs compared to surface has only been estimated using short reads to a less contiguous reference genome (Warren *et al*. 2021). To evaluate moderately-sized SVs (50 to 10,000 bp) among these cave morphs compared to the surface genome, we estimated the presence of deletions, insertions, and contractions and expansions of repeats using Assemblytics (Nattestad and Schatz 2016). The number of insertions and deletions shows comparable total bases affected across these cave morphs relative to surface regardless of their sequence length distribution (500-10,000 bp Fig. 3;50-500 bp Suppl. Fig. 3; Suppl. Table 10). The average total size of detected cave morph deletions and insertions was 19.4 MB (1.42% of the genome size) when aligned to the surface genome (Suppl. Table 10). SV divergences unique to each cave morph were also evident. For example, the Tinaja morph had the highest total sequence size for all insertions (12.9 MB), followed by Pachón (12.8 Mb) and Molino (10.8 Mb) (Suppl. Table 10). Similarly for all deletions, Tinaja was highest (7.54 MB), then Pachón (7.10 Mb), and Molino (7.08Mb) (Suppl. Table 10). Interestingly, total inserted sequence by size distribution changes cave morph order with Pachón (9.9 Mb) having the largest amount for the 500 to 10,000 bp range followed by Tinaja (9.7 Mb) (Suppl. Table 10). When evaluating the cave morph repeat landscape, we find an even larger percentage of the genome impacted relative to insertions and deletions, with the average repeat expansion or contraction being 3.05 and 2.91% of the total genome, respectively (Suppl. Table 10). These broadly classified SV and repeat results highlight their differences vary by category and cave morph origin, which may be the result of numerous factors, including assembly completeness and accuracy, mixed haplotype assembly architecture, and the diversified origins of each cave morph reference (Herman *et al*. 2018).

**Figure 3.**
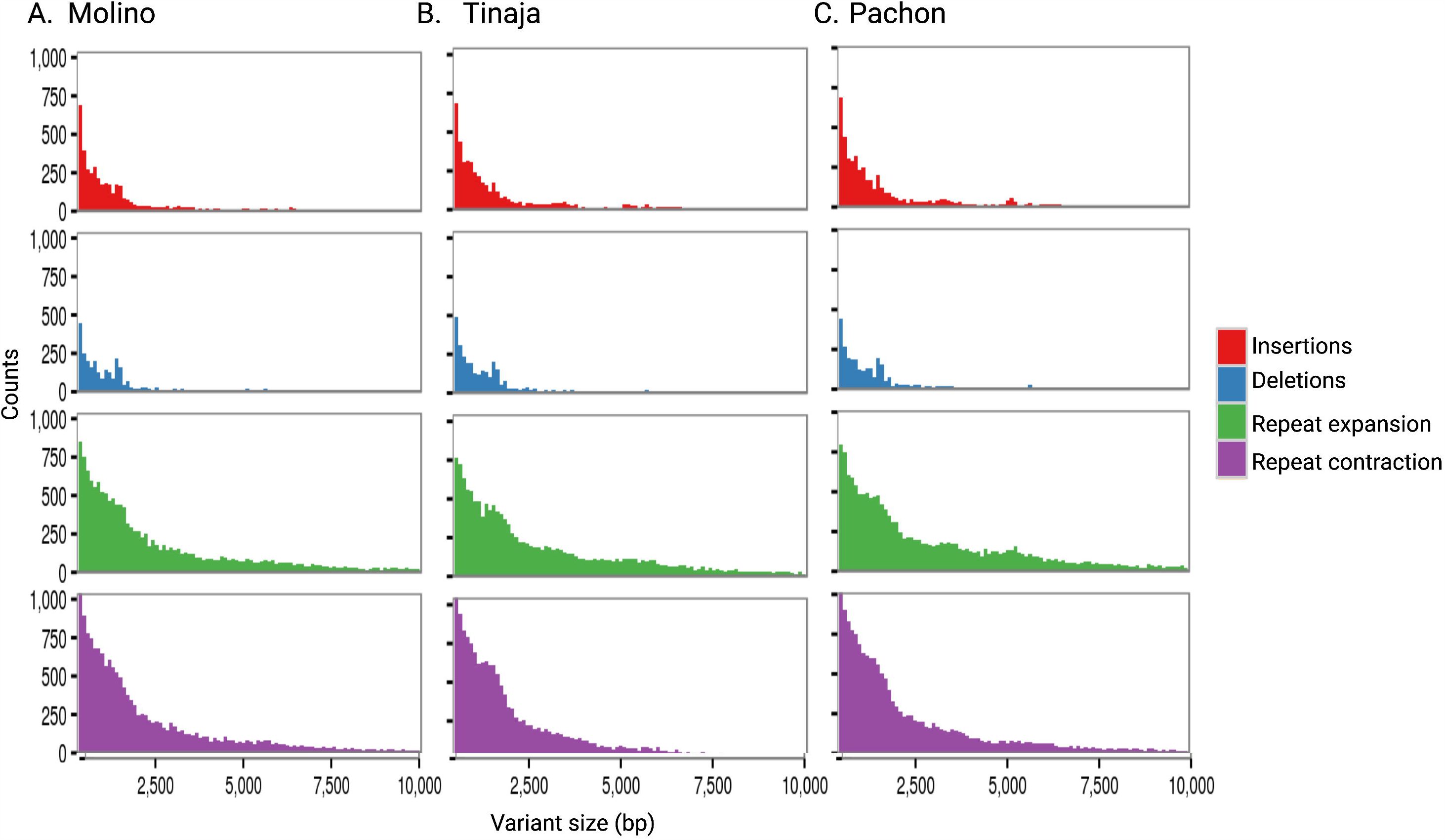
Genomic structural variation among *A. mexicanus* cave morph assemblies when compared to surface. Cave morph specific distribution by counts for A. Molino, B. Tinaja, and C. Pachón for the size distribution of 500 to 10,000 bp.

### Study conclusions

The surface-to-cave genomic transitions that occurred in *A. mexicanus* establish a unique model for the study of natural polygenic trait adaptation. Here, we provide initial evidence that these more complete genomes will substantially advance our capability to resolve these signatures of genetic adaption. The availability of nearly complete genome copies of a surface and the independently evolved cave morphs will drive future reevaluations of all types of segregating single nucleotide variants and SVs in a pangenome-dependent manner (Siren *et al*. 2021).

## Acknowledgments

We thank the Stowers aquatics group for fish care and help with shipments of fish samples. AK, SEM, NR are supported by NIH 1R01GM127872-01. This work was also supported by NIH R24OD011198 to WCW, AK, NR. NSF EDGE Award 1923372 to NR, SEM, and NSF IOS-1933428 to SEM, NR. Wellcome Trust, award number WT222155/Z/20/Z to LH, DO, and FM. Some computation for this work was performed on the high-performance computing infrastructure provided by Research Computing Support Services and in part by the National Science Foundation under grant number CNS-1429294 at the University of Missouri, Columbia MO. The Minnesota Supercomputing Institute (MSI) at the University of Minnesota provided resources that contributed to the research results reported within this paper.

## Data availability

All raw and processed data for this study are available by querying NCBI BioProject accession numbers PRJNA807270, PRJNA819394, and PRJNA819399. In addition, each assembly is available under GenBank numbers GCA_023375975.1, GCA_023375835.1, and GCA_023375845.1. The full availability of orthofinder results are attainable on figshare.

## Code availability

Scripts used for this study are available at the following GitHub repositories:

https://github.com/esrice/hic-pipeline

https://github.com/WarrenLab/purge-haplotigs-nf

https://github.com/WarrenLab/agptools

## Author notes

### Competing interests

The authors declare no competing interests.

## Notes

### Competing Interest Statement

The authors have declared no competing interest.

### Summary of Updates

The author Fergal Smith is changed to Fergal Martin

## Literature cited

Adrian-Kalchhauser, I., A. Blomberg, T. Larsson, Z. Musilova, C. R. Peart et al., 2020 The round goby genome provides insights into mechanisms that may facilitate biological invasions. BMC Biol 18: 11.

Aken, B. L., S. Ayling, D. Barrell, L. Clarke, V. Curwen et al., 2016 The Ensembl gene annotation system. Database 2016.

Albalat, R., and C. Canestro, 2016 Evolution by gene loss. Nat Rev Genet 17: 379–391.

Aspiras, A. C., N. Rohner, B. Martineau, R. L. Borowsky and C. J. Tabin, 2015 Melanocortin 4 receptor mutations contribute to the adaptation of cavefish to nutrient-poor conditions. Proc Natl Acad Sci U S A 112: 9668–9673.

Bilandžija, H., Hollifield, B., Steck, M., Meng, G., Ng, M., Koch, A.D., Gračan, R.,Ćetković, H., Porter, M.L., Renner, K.J., Jeffery, W.R., 2019 Phenotypic plasticity as an important mechanism of cave colonization and adaptation in Astyanax cavefish. bioRxiv.

Braasch, I., A. R. Gehrke, J. J. Smith, K. Kawasaki, T. Manousaki et al., 2016 The spotted gar genome illuminates vertebrate evolution and facilitates human-teleost comparisons. Nat Genet 48: 427–437.

Cheng, H., G. T. Concepcion, X. Feng, H. Zhang and H. Li, 2021 Haplotype-resolved de novo assembly using phased assembly graphs with hifiasm. Nat Methods 18: 170–175.

Chernyavskaya, Y., X. Zhang, J. Liu and J. Blackburn, 2022 Long-read sequencing of the zebrafish genome reorganizes genomic architecture. BMC Genomics 23: 116.

Du, K., M. Pippel, S. Kneitz, R. Feron, I. da Cruz et al., 2022 Genome biology of the darkedged splitfin, Girardinichthys multiradiatus, and the evolution of sex chromosomes and placentation. Genome Res 32: 583–594.

Duboue, E. R., A. C. Keene and R. L. Borowsky, 2011 Evolutionary convergence on sleep loss in cavefish populations. Curr Biol 21: 671–676.

Durand, N. C., J. T. Robinson, M. S. Shamim, I. Machol, J. P. Mesirov et al., 2016 Juicebox Provides a Visualization System for Hi-C Contact Maps with Unlimited Zoom. Cell Syst 3: 99–101.

Emms, D. M., and S. Kelly, 2019 OrthoFinder: phylogenetic orthology inference for comparative genomics. Genome Biol 20: 238.

Ghurye, J., and M. Pop, 2019 Modern technologies and algorithms for scaffolding assembled genomes. PLoS Comput Biol 15: e1006994.

Ghurye, J., A. Rhie, B. P. Walenz, A. Schmitt, S. Selvaraj et al., 2019 Integrating Hi-C links with assembly graphs for chromosome-scale assembly. PLoS Comput Biol 15: e1007273.

Guan, D., S. A. McCarthy, J. Wood, K. Howe, Y. Wang et al., 2020 Identifying and removing haplotypic duplication in primary genome assemblies. Bioinformatics 36: 2896–2898.

Harrison, P. M., 2021 Computational Methods for Pseudogene Annotation Based on Sequence Homology. Methods Mol Biol 2324: 35–48.

Herman, A., Y. Brandvain, J. Weagley, W. R. Jeffery, A. C. Keene et al., 2018 The role of gene flow in rapid and repeated evolution of cave-related traits in Mexican tetra, Astyanax mexicanus. Mol Ecol 27: 4397–4416.

Imarazene, B., K. Du, S. Beille, E. Jouanno, R. Feron et al., 2021 A supernumerary “Bsex” chromosome drives male sex determination in the Pachon cavefish, Astyanax mexicanus. Curr Biol 31: 4800–4809 e4809.

J., D., 2020 AGAT: Another Gff Analysis Toolkit to handle annotations in any GTF/GFF format., pp. in Zenodo.

Jarvis, E. D., G. Formenti, A. Rhie, A. Guarracino, C. Yang et al., 2022 Semi-automated assembly of high-quality diploid human reference genomes. Nature 611: 519–531.

Kurtz, S., A. Phillippy, A. L. Delcher, M. Smoot, M. Shumway et al., 2004 Versatile and open software for comparing large genomes. Genome Biol 5: R12.

Li, H., and R. Durbin, 2009 Fast and accurate short read alignment with Burrows-Wheeler transform. Bioinformatics 25: 1754–1760.

Lovell, J. T., A. Sreedasyam, M. E. Schranz, M. Wilson, J. W. Carlson et al., 2022 GENESPACE tracks regions of interest and gene copy number variation across multiple genomes. Elife 11.

McGaugh, S. E., J. E. Kowalko, E. Duboue, P. Lewis, T. A. Franz-Odendaal et al., 2020 Dark world rises: The emergence of cavefish as a model for the study of evolution, development, behavior, and disease. J Exp Zool B Mol Dev Evol 334: 397–404.

Moore, B., M. Herrera, E. Gairin, C. Li, S. Miura et al., 2023 The chromosome-scale genome assembly of the yellowtail clownfish Amphiprion clarkii provides insights into the melanic pigmentation of anemonefish. G3 (Bethesda) 13.

Moran R.L. R. E.J., Ornelas-García, C.P., Gross, J.B., Donny, A., Wiese, J., Keene, A.C., Kowalko, J.E., Rohner, N., McGaugh, S.E., 2022 Selection-driven trait loss in independently evolved cavefish populations. bioRxiv.

Nattestad, M., and M. C. Schatz, 2016 Assemblytics: a web analytics tool for the detection of variants from an assembly. Bioinformatics 32: 3021–3023.

Ojha, A., and M. Watve, 2018 Blind fish: An eye opener. Evol Med Public Health 2018: 186–189.

Pruitt, K. D., G. R. Brown, S. M. Hiatt, F. Thibaud-Nissen, A. Astashyn et al., 2014 RefSeq: an update on mammalian reference sequences. Nucleic Acids Res 42: D756–763.

Rautiainen, M., S. Nurk, B. P. Walenz, G. A. Logsdon, D. Porubsky et al., 2023 Telomere-to-telomere assembly of diploid chromosomes with Verkko. Nat Biotechnol.

Rhie, A., S. A. McCarthy, O. Fedrigo, J. Damas, G. Formenti et al., 2021 Towards complete and error-free genome assemblies of all vertebrate species. Nature 592: 737–746.

Roberts, M. B., D. T. Schultz, R. Gatins, M. Escalona and G. Bernardi, 2023 Chromosome-level genome of the three-spot damselfish, Dascyllus trimaculatus. G3 (Bethesda) 13.

Robinson, J. T., D. Turner, N. C. Durand, H. Thorvaldsdottir, J. P. Mesirov et al., 2018 Juicebox.js Provides a Cloud-Based Visualization System for Hi-C Data. Cell Syst 6: 256–258 e251.

Rohner, N., 2018 Cavefish as an evolutionary mutant model system for human disease. Dev Biol 441: 355–357.

Ruan, J., and H. Li, 2020 Fast and accurate long-read assembly with wtdbg2. Nat Methods 17: 155–158.

Simao, F. A., R. M. Waterhouse, P. Ioannidis, E. V. Kriventseva and E. M. Zdobnov, 2015 BUSCO: assessing genome assembly and annotation completeness with single-copy orthologs. Bioinformatics 31: 3210–3212.

Siren, J., J. Monlong, X. Chang, A. M. Novak, J. M. Eizenga et al., 2021 Pangenomics enables genotyping of known structural variants in 5202 diverse genomes. Science 374: abg8871.

Stockdale, W. T., M. E. Lemieux, A. C. Killen, J. Zhao, Z. Hu et al., 2018 Heart Regeneration in the Mexican Cavefish. Cell Rep 25: 1997–2007 e1997.

Volff, J. N., 2005 Genome evolution and biodiversity in teleost fish. Heredity (Edinb) 94: 280–294.

Warren, W. C., T. E. Boggs, R. Borowsky, B. M. Carlson, E. Ferrufino et al., 2021 A chromosome-level genome of Astyanax mexicanus surface fish for comparing population-specific genetic differences contributing to trait evolution. Nat Commun 12: 1447.

Warrenlab., 2022 Nextflow workflow for purging haplotigs from a genome assembly.

