## Supplemental Figures for "Astyanax mexicanus surface and cavefish chromosome-scale assemblies for trait variation discovery"

A.
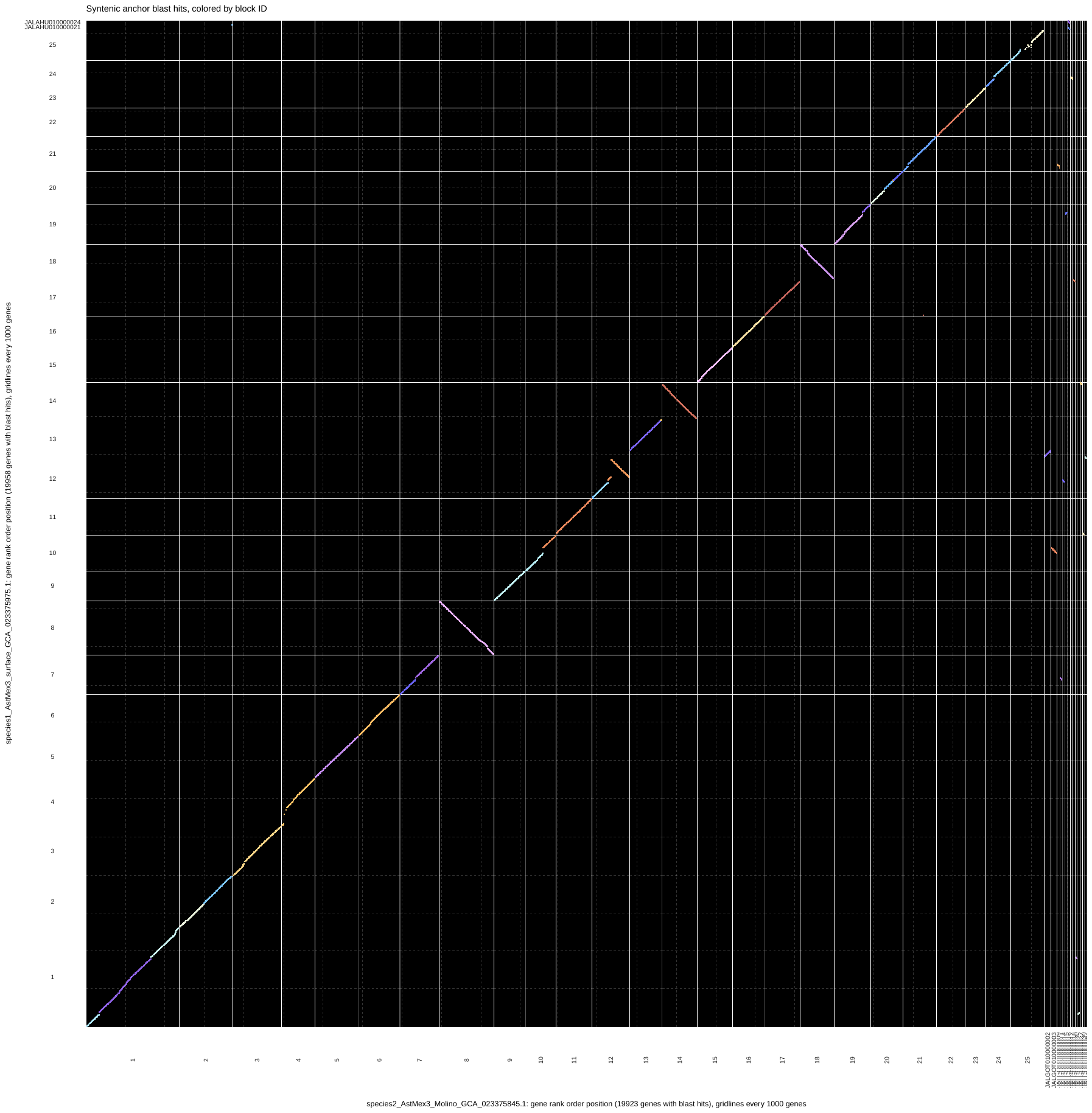

Chromosomes (Surface)

Chromosomes (Molino)

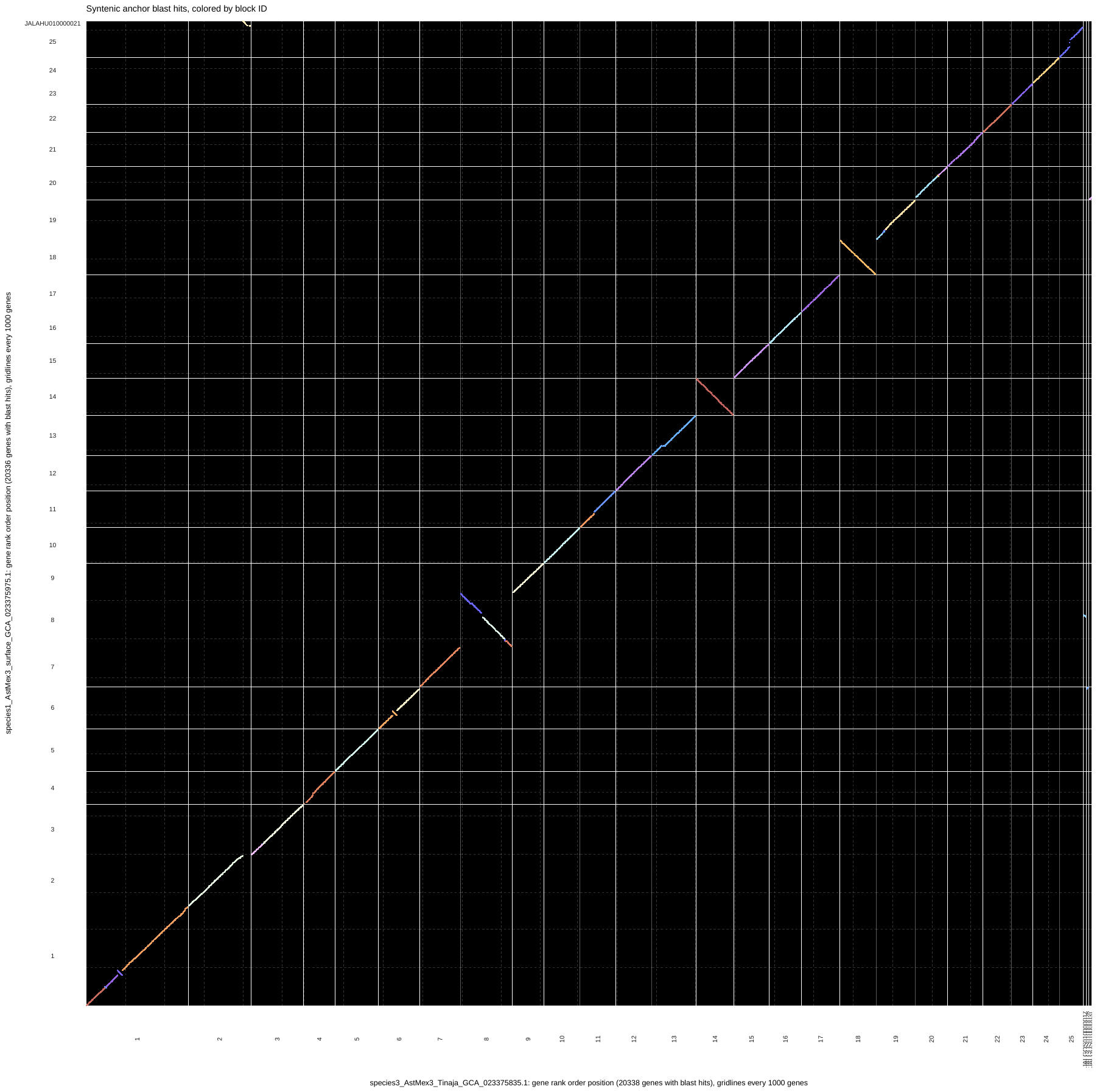
.

B.

Chromosomes (Surface)

Chromosomes (Tinaja)

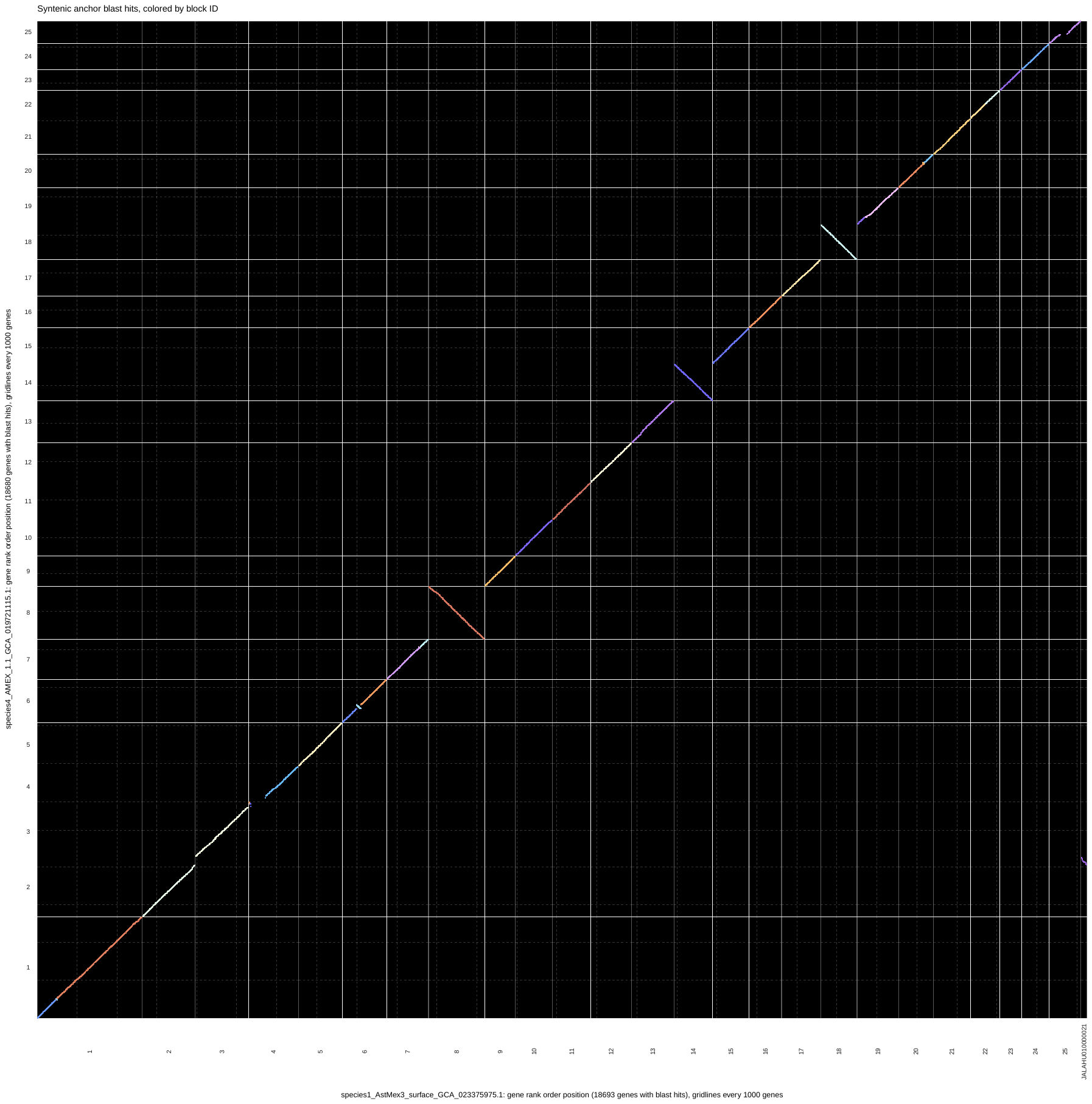

C.

Chromosomes (Surface)

Chromosomes (Pachon)

Figure 1. Chromosomal synteny of all cave morphs when compared to the surface genome using Genespace for A. Molino (19,923 blast hits), B. Tinaja (20,338 blast hits), and C. Pachon (18,693 blast hits). Axes gridlines are every 1,000 genes.

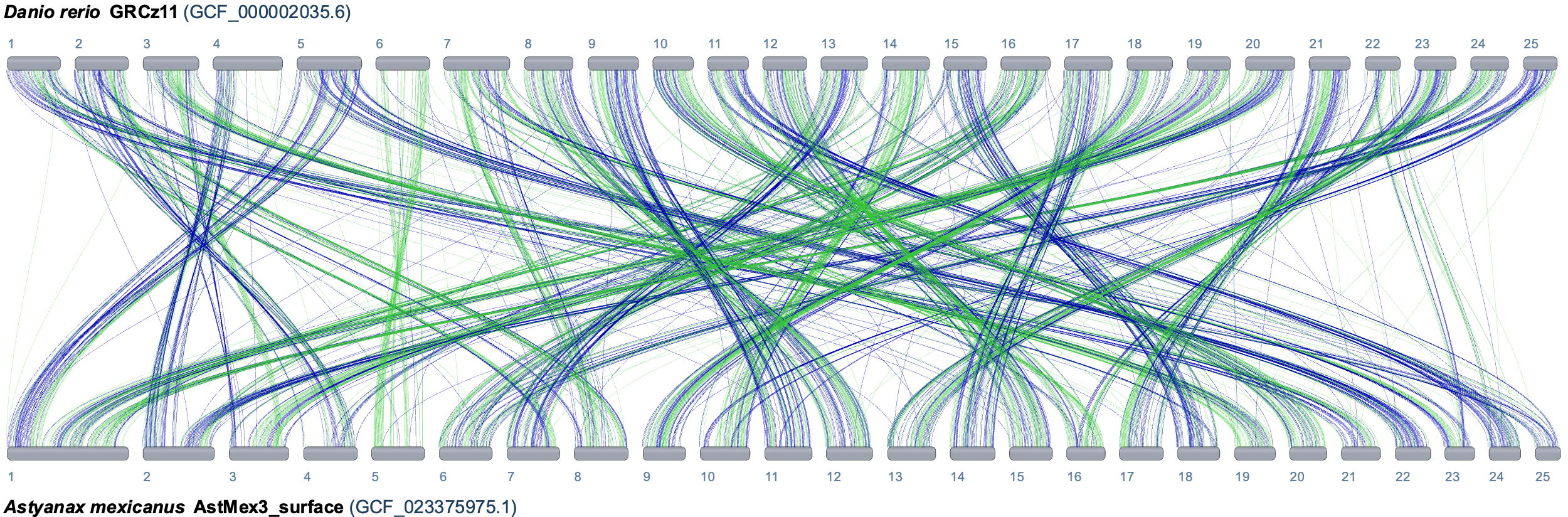

Figure 2. Whole genome alignments of zebrafish chromosomes to the *A. mexicanus* genome using the NCBI comparative genomes browser.

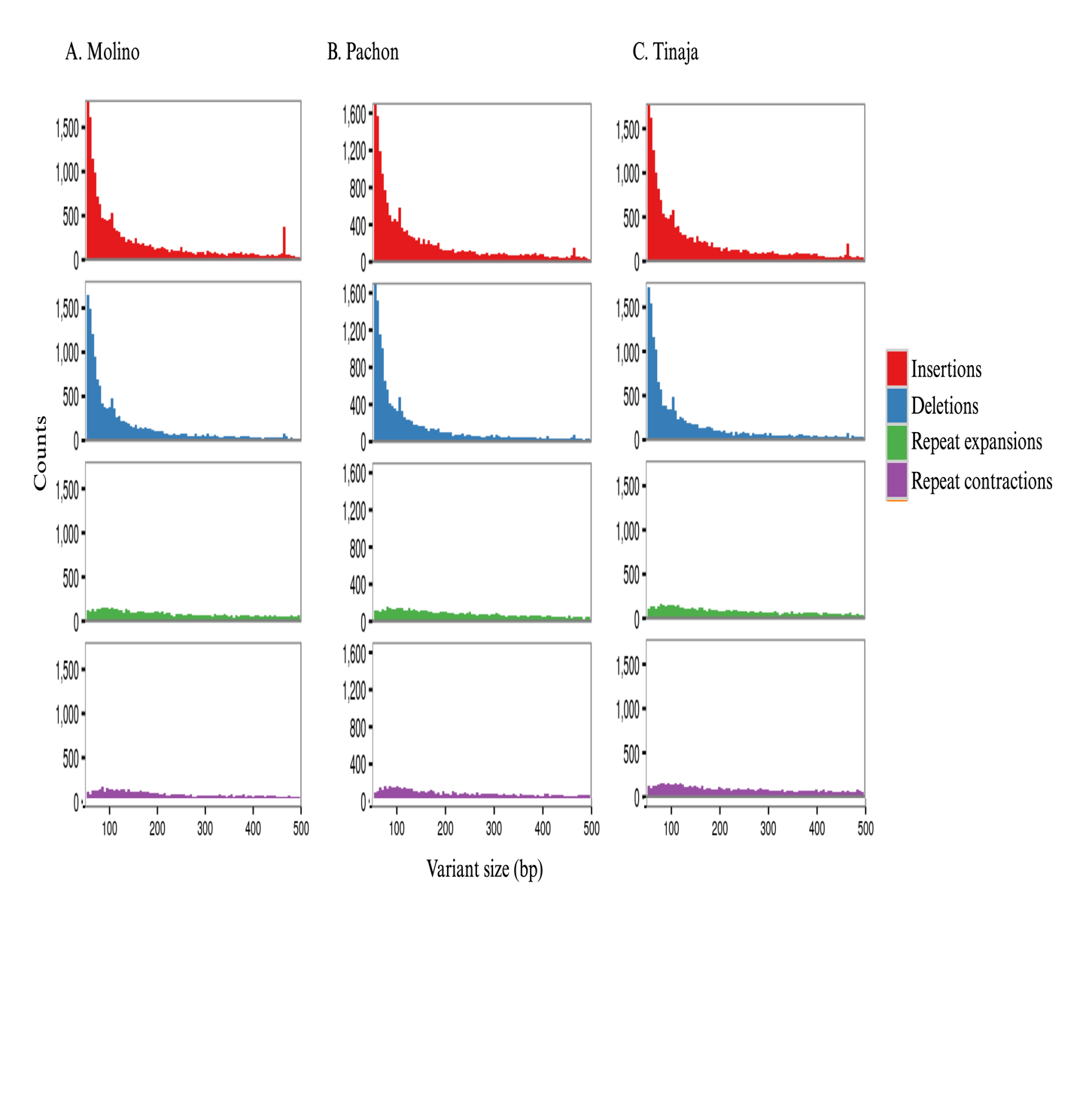

Figure 3. The distribution of SVs for all cave morphs compared to surface fish using the Assemblytics output. The size range of 50 to 500bp is show here for Molino, Pachon, and Tinaja.
